# NLRP1B and NLRP3 control the host response following colonization with the commensal protist *Tritrichomonas musculis*

**DOI:** 10.1101/2021.08.14.455483

**Authors:** Pailin Chiaranunt, Kyle Burrows, Louis Ngai, Eric Y. Cao, Helen Liang, Catherine J. Streutker, Stephen E. Girardin, Arthur Mortha

## Abstract

Commensal intestinal protozoa, unlike their pathogenic relatives, are neglected members of the mammalian microbiome. These microbes have a significant impact on the host’s intestinal immune homeostasis, typically by elevating anti-microbial host defense. *Tritrichomonas musculis* (*T. mu*), a protozoan gut commensal, strengthens the intestinal host defense against enteric Salmonella infections through *Asc*- and *Il1r1*-dependent Th1 and Th17 cell activation. However, the underlying inflammasomes mediating this effect remain unknown. Here, we report that colonization with *T. mu* results in an increase in luminal extracellular ATP, elevated levels of IL-1β, and increased numbers of IL-18 receptor-expressing Th1 and Th17 cells in the colon. Mice deficient in either *Nlrp1b* or *Nlrp3* failed to display these protozoan-driven immune changes and lost resistance to enteric Salmonella infections even in the presence of *T. mu*. These findings demonstrate that *T. mu*-mediated host protection requires sensors of extra and intracellular ATP to confer resistance to enteric Salmonella infections.

**KEY POINTS:**

- Intestinal colonization with the commensal *Tritrichomonas musculis* elevates luminal ATP levels
- NLRP1B and NLRP3 activation is required for *Tritrichomonas musculis-*driven Th cell response.

## Introduction

The mammalian intestinal microbiota is a rich multi-kingdom ecosystem, comprising of bacteria, viruses, protozoa, fungi, and archaea that modulate host immunity. A diverse multi-kingdom microbiota has been suggested to increase host fitness and improve outcomes in enteric infections (1, 2). Recent work has begun to elucidate the roles that these mostly neglected non-bacterial commensal microorganisms have in this community, revealing key mechanisms of innate and adaptive immune responses to the mycobiota (3-5) and enteric virome (6, 7). The central sensory elements of these microbes include Toll-like receptors (TLRs) and Nod-like receptors (NLRs), C-type lectin receptors (CLRs), and RIG-I-like receptors (RLRs), which are required for the host recognition of bacteria, fungi, and viruses, respectively (8-10). Much of the work investigating the role of sensory systems for helminths and pathogenic protozoa (e.g. *Toxoplasma* spp., *Giardia* spp., or *Cryptosporidium* spp.) revealed a combinatorial requirement of TLRs, NLRs, and taste receptors for effective host sensing and parasite defense (11, 12). However, other less prominent members of the protozoa kingdom are considered “commensal-like” (13, 14). For example, *Dientamoeba fragilis* (*D. fragilis*), a member of the order Trichomonadida, often shows asymptomatic colonization in humans (15). *Tritrichomonas* spp., including the recently identified *Tritrichomonas musculis* (*T. mu*), *Tritirichomonas muris* (*T. muris*), and *Tritrichomonas raineri* (*T. raineri*), are commensal protozoa found in wild animals and laboratory mice at certain facilities (12, 16-18) (Mortha A., unpublished observation). Unlike pathogenic protozoa, these gut-dwelling Tritrichomonads are not cleared by the immune system, instead resulting in chronic, asymptomatic colonization in mice following horizontal or vertical transmission (17). Nevertheless, their engraftment into the gut microbiome induces a strong innate and adaptive immune response in the host (11, 12, 16-18). All three *Tritrichomonas* spp. were found to activate small intestinal group 2 innate lymphoid cells (ILC2) via Tuft cell-derived interleukin(IL)-25 (11, 12, 17, 18). Interestingly, colonization of mice with *T. musculis* or *T. muris* additionally resulted in prominent and persistent induction and expansion of colonic Th1 and Th17 cells (16, 17). The elevated differentiation of interferon (IFN)-γ- and IL-17A-producing helper T (Th) cells exacerbated T cell-driven autoimmunity and sporadic colorectal cancer, while conferring resistance to enteric infections by *Salmonella typhimurium* (*S. typhimurium*) *(16, 17)*. The Th1 and Th17 responses required *Asc* (apoptosis-associated speck-like protein containing a CARD), *Il18*, and *Il1r1*, suggesting that ASC-associated NLRs may be required for the detection of *T. mu* in the gut *(17)*.

Here, we demonstrate that colonization with *T. mu* results in elevated levels of luminal extracellular adenosine 3’-triphosphate (ATP), accompanied by increased concentrations of colonic IL-1β. Colonic Th1 and Th17 cells in *T. mu*-colonized mice expressed the IL-18 receptor (IL-18R) and protected the host from *S. typhimurium*-induced pathology and tissue dissemination. *T. mu*-driven activation of Th1 and Th17 cells was dependent on *Nlrp1b* and in parts on *Nlrp3*. In line with these findings, *Nlrp1b*^*-/-*^ and *Nlrp3*^*-/-*^ mice displayed impaired protection against enteric *Salmonella* infection despite the presence of *T. mu*. These findings demonstrate that *T. mu*-mediated host immune modulation requires the NLRP1B and NLRP3 inflammasomes to confer host protection against Salmonella infections.

## Materials and Methods

### Mice

C57BL/6, B6.129S6-*Nlrp1b*^*tm1Bhk*^/J (*Nlrp1b*^-/-^), B6.129S6-*Nlrp3*^*tm1Bhk*^/J (*Nlrp3*^-/-^), and B6.129P2-*P2rx7*^*tm1Gab*^/J (*P2xr7*^*-/-*^) mice were purchased from Jackson Laboratory and subsequently bred in-house under specific pathogen-free conditions at the University of Toronto, Division of Comparative Medicine. All experiments were conducted using age- and sex-matched littermates and with approval by the animal care committee, University of Toronto.

### Purification and colonization of *Tritrichomonas musculis*

Cecal contents of *T. mu*^+^ mice were collected, resuspended in PBS, filtered through a 70 µm cell strainer, and spun for 10min at 600 x *g*. The resulting pellet was put through a 40/80% Percoll gradient centrifugation. The *T. mu*-enriched interphase was collected. Protozoa were then resuspended in PBS and double sorted into PBS based on size, granularity, and violet autofluorescence on a FACS ARIA II. Two million *T. mu* were orally gavaged into mice immediately after the sort.

### Isolation of colonic lamina propria and mesenteric lymph node leukocytes

Colonic lamina propria (LP) cells were isolated as previously described(19). Briefly, colons were washed in HBSS plus 5 mM EDTA and 10 mM HEPES to strip the epithelium. Tissues were then minced and shaken at 37°C for 20 min in digestion buffer (HBSS with calcium and magnesium, supplemented with 10 mM HEPES, 4% FBS, penicillin-streptomycin (Sigma Aldrich), 0.5 mg/mL DNase I (Sigma Aldrich), and 0.5 mg/mL Collagenase (Sigma Aldrich)). Supernatants were collected and enriched for leukocytes using a 40/80% Percoll gradient, after which cells are ready for downstream use. MLN cells were mashed through a 70 µm cell strainer and resuspended in FACS buffer (PBS with 2% FBS and 5 mM EDTA).

### Salmonella infection and pathological assessment

Groups of age- and sex-matched littermates were orally gavaged with either PBS or 2 × 10^6^ purified *T. mu*. 3 weeks later, mice were orally gavaged with streptomycin and infected with *Salmonella typhimurium* as previously described (20). Mice were euthanized 48h later, and cecal weight was recorded. Cecal pieces were fixed, embedded in paraffin, sectioned, and stained with hematoxylin and eosin (H&E) according to standard procedures. Pathological evaluation was performed in a blinded fashion by a pathologist and scored as previously described (20). Colony forming units (CFUs) of *S. typhimurium* in feces, colon, cecum, mLN, liver, and spleen were measured on MacConkey agar plates containing 50 µg/mL streptomycin.

### Flow cytometry

For intracellular staining, cells were first stimulated for 4 h with PMA, ionomycin, and protein transport inhibitor cocktail containing Brefeldin A and Monensin (eBioscience). Cells were then incubated on ice for 20 min with Fc block (CD16/CD32; eBioscience), surface markers, and Fixable Viability Dye eFluor™ 506 (eBioscience). Cells were fixed and permeabilized using the BD Cytofix/Cytoperm Kit, followed by cytokine stains, then re-fixed and permeabilized using the eBioscience Foxp3/Transcription Factor Staining Buffer Set, followed by transcription factor stains. Samples were analyzed on an LSR Fortessa X-20 (BD).

For surface staining, the following anti-mouse Abs were used: TCRβ (H57-597; eBioscience), CD4 (GK1.5; BioLegend), CD45 (30-F11; BioLegend), CD218a (IL-18Ra) (P3TUNYA; eBioscience), ST2 (RMST2-2; eBioscience), CD11b (M1/70; BioLegend), Ly6c (HK1.4; eBioscience), CD64 (X54-5/7.1; BioLegend), and MHCII (I-A/I-E) (M5/114.15.2; eBioscience). Intracellular markers include anti-mouse IFN-γ (XMG1.2; eBioscience), TNFα (MP6-XT22; eBioscience), IL-10 (JESS-16E3; BioLegend), IL-17A (TC11-18H10.1; BioLegend), and FOXP3 (MF-14; BioLegend). CD4^+^ T cells were gated as Live CD45^+^ TCRβ^+^ CD4^+^. Immature macrophages were gated as Live CD45^+^ CD64^+^ CD11b^+^ Ly6c^hi^ MHCII^lo^.

### ELISA

Proximal colon explants were weighed, washed in RPMI supplemented with 50 ug/mL gentamicin (Gibco) for 30 minutes at room temperature, then cultured in 500 uL complete RPMI plus 5% FBS (Gibco), 50 ug/mL gentamicin (Sigma Aldrich), and penicillin-streptomycin (Sigma Aldrich) for 18-24 h. Supernatants were collected and used for cytokine measurement with IL-18 (Invitrogen; sensitivity 19.0 pg/mL), IL-1β (Invitrogen; sensitivity 8 pg/mL), and IL-33 (Invitrogen; sensitivity 25 pg/mL) ELISA kits according to the manufacturer’s instructions.

### Luminal ATP measurement

Fecal samples were collected, homogenized in PBS plus 0.01% NaN3 using the Omni Bead Ruptor 24, and centrifuged twice (800 x *g* followed by 10000 x *g*) to remove debris and microbes. Supernatants were filtered through a 0.2 µm filter, then analyzed for ATP levels using the ENLITEN ATP Assay System Bioluminescence Detection Kit (Promega) according to the manufacturer’s instructions.

### Statistical analyses

All data are shown as mean ± SEM. Statistical tests were performed using GraphPad Prism as detailed in figure legends.

## Results and Discussion

### Colonization with *T. mu* increases luminal extracellular ATP levels and IL-1β release in the colon

Activation of colonic Th1 and Th17 cells following colonization with the commensal protist *T. mu* requires ASC, IL-18, and IL-1R (17). Extracellular ATP is a ubiquitously produced metabolite and danger associated molecular pattern (DAMP) that facilitates the production of active IL-18 and IL-1β in an ASC-dependent manner (21). Luminal ATP levels have previously been implicated in the differentiation of Th17 cells and may derive from commensal bacteria or dying host cells (22). To determine whether colonization of wild-type (WT) mice with *T. mu* affects the extracellular ATP concentrations in the gut lumen, ATP levels were measured in fecal samples using a luciferase-based luminescence assay. Following colonization for 21 days, mice carrying *T. mu* displayed significantly elevated concentrations of extracellular ATP in the gut lumen (Figure 1A). In line with previous reports, increased levels of extracellular ATP resulted in higher IL-1β concentrations in colonic explants (Figure 1B) (22). Interestingly, levels of IL-18 and the type 2 immunity-associated alarmin IL-33 did not show a significant increase following colonization by *T. mu*, despite reports of the latter exhibiting ATP-dependent activation at other mucosal surfaces (Figure 1C) (23). These results indicate that colonization with the protozoan commensal *T. mu* results in the local activation of the innate immune system.

**Figure 1.**
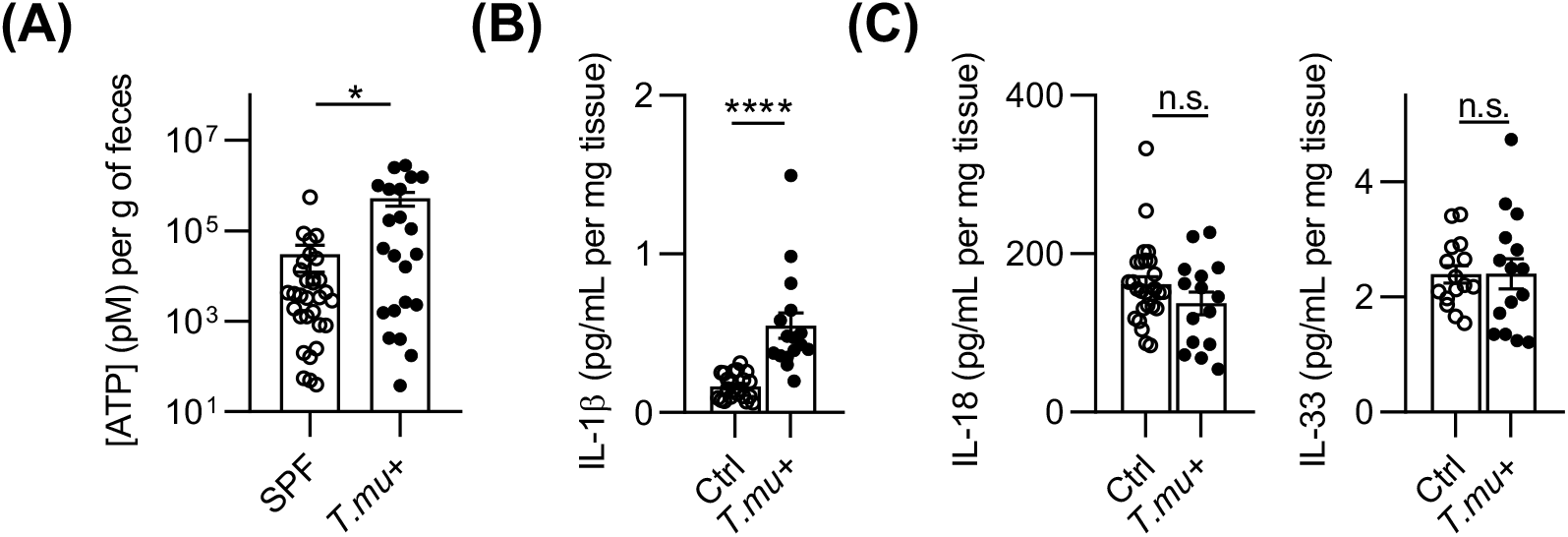
Colonization with *T. mu* increases luminal extracellular ATP levels and IL-1β release in the colon. (A-C) C57BL/6 mice were either left untreated or orally gavaged with 2 × 10^6^ purified *T. mu*. Analysis was performed 3 weeks post-gavage. (A) Levels of extracellular ATP were measured in the fecal supernatants of freshly isolated fecal pellets using the Promega ENLITEN ATP Assay as per the manufacturer’s instructions. (B and C) Colonic explants were placed in RPMI + gentamicin for 30 min, then incubated overnight at 37°C in 500 uL RPMI supplemented with gentamicin, penicillin/streptomycin, and 5% FBS. Supernatants were collected for IL-1β, IL-18, or IL-33 ELISA analysis. Data shown is representative of at least three independent experiments with at least three mice per group per experiment. Data show mean ± SEM. Unpaired student’s t test was performed; *p < 0.05, ****p < 0.0001; n.s., not significant.

### The protozoan commensal *T. mu* induces the accumulation of IL-18R^+^ Th1 and Th17 cells in the colonic lamina propria

To determine the effects of *T. mu* colonization on Th cell activation, we analyzed IFN-γ and IL-17A production in CD4^+^ T cells in the colonic lamina propria. Mice colonized with *T. mu* displayed significantly increased percentage and absolute number of IFN-γ^+^ Th1 cells and IL-17A^+^ Th17 cells (Figure 2A). This induction of Th1 and Th17 cells required viable protist, as oral gavage of heat-killed *T. mu* did not result in an elevation of these cells in the colonic lamina propria (Supplementary Figure 1A). In line with previous findings, colonization with *T. mu* increased the number of IL-18R-expressing CD4^+^ Th cells, including IFN-γ^+^ Th1 cells and IL-17A^+^ Th17 cells (Figure 2B and 2C) (17). ST2, the receptor of IL-33 is involved in the induction of Th2 and regulatory T cells and has been shown to contribute to the anti-viral Th1 response during the acute effector phase (24, 25). However, IFN-γ^+^ Th1 cells in mice colonized with *T. mu* did not express increased levels of ST2, suggesting that their polarization does not depend on IL-33 (Figure 2D). The expression of IL-18R on human and murine cells, including intestinal epithelial cells, has previously been reported to require TNFα, which can be produced by activated Th1 cells (26, 27). To determine whether IL-18R expression on Th cells would correlate with elevated TNFα release by colonic Th cells, we assessed TNFα production of these cells using flow cytometry. Th cells did not release increased levels of TNFα after colonization with *T. mu*, suggesting alternative sources of TNFα as a possible reason for the increase in IL-18R-expressing Th cells (Supplementary Figure 1B). We previously demonstrated changes in the myeloid immune landscape following colonization with *T. mu*. These changes included an increase in colonic dendritic cells (DCs) migrating to the mesenteric lymph nodes and an accumulation of Ly6C^+^ TNFα^+^ immature macrophages in the lamina propria of *T. mu*-colonized mice (17). Macrophage activation by extracellular ATP uses a P2X7R-dependent mechanism (21, 22). To investigate whether the accumulation of immature macrophages in the colon of *T. mu*-colonized mice required ATP-mediated activation of P2X7R, infiltrating Ly6C^+^ macrophage counts were quantified following colonization with *T. mu* in *P2xr7*^*-/-*^ mice and age- and sex-matched littermate controls. In line with our previous report, WT littermate control mice showed a significant increase in immature macrophages in the presence of *T. mu*, while the colons of *P2xr7*^-/-^ mice remained unchanged in macrophage numbers after colonization with the protist (Supplementary Figure 1C). These data suggest that *T. mu* colonization facilitates the P2X7R-dependent accumulation of immature macrophages, a potent source for TNFα, followed by the release of IL-1β and the differentiation of Th1 and Th17 cells in an IL-18R-dependent manner.

**Figure 2.**
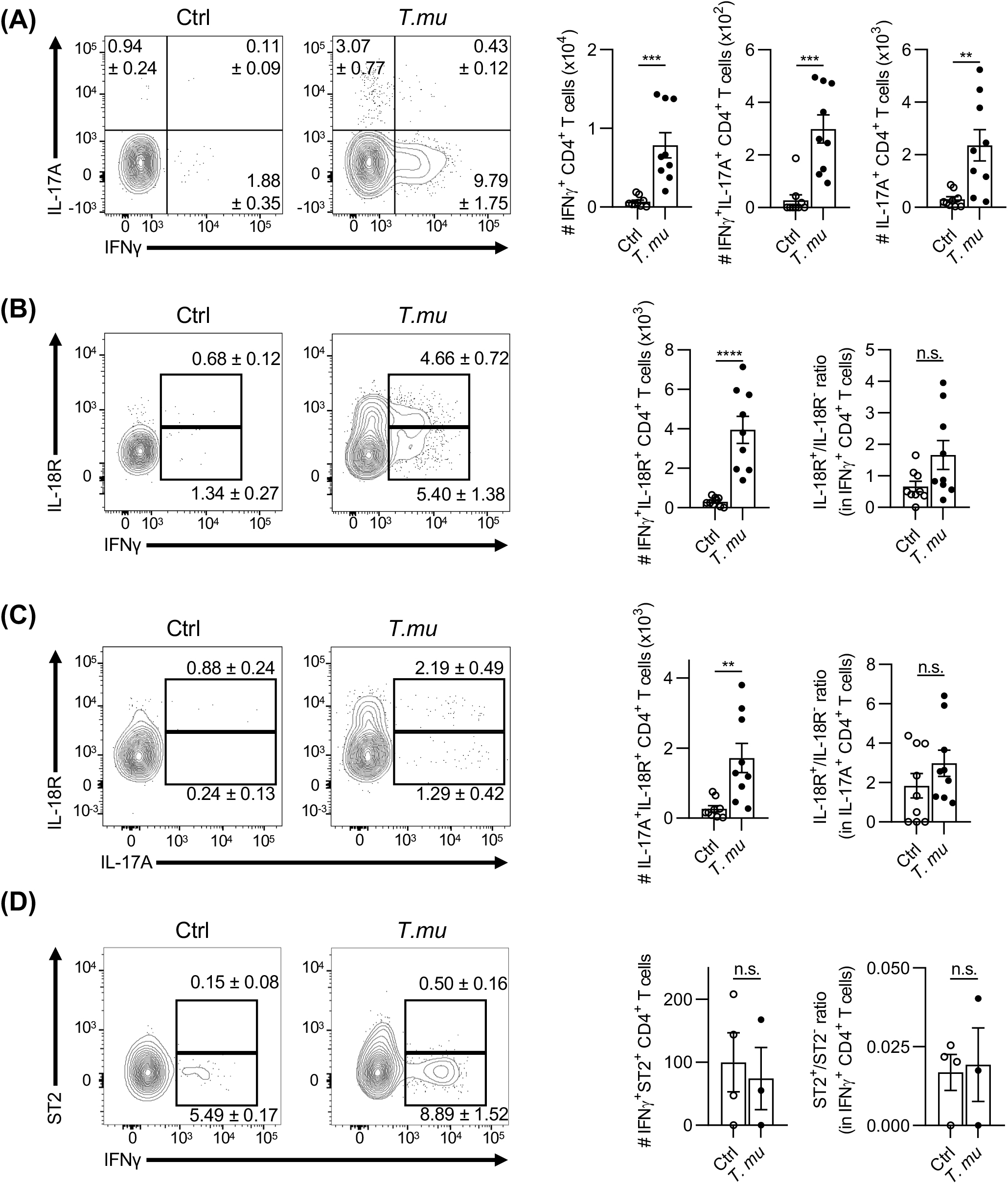
The protozoan commensal *T. mu* induces accumulation of IL-18R^+^ Th1 and Th17 cells in the lamina propria. C57BL/6 mice were either left untreated or orally gavaged with 2 × 10^6^ purified *T. mu*. Colons were harvested 3 weeks later. Colonic lamina propria leukocytes were isolated, stimulated for 4 h with PMA and ionomycin in the presence of protein transport inhibitor cocktail, and then stained and analyzed via flow cytometry for (A) IFN-γ and IL-17A production by CD4^+^ T cells, IL-18R expression in (B) IFNγ-producing CD4^+^ T cells and (C) IL-17A-producing CD4^+^ T cells, or (D) ST2 expression in IFNγ-producing CD4^+^ T cells. Bar graphs show absolute numbers of cell populations represented in respective flow cytometry plots. Data shown is representative of at least three independent experiments with at least three mice per group per experiment. Data show mean ± SEM. Student’s t test was performed; *p < 0.05, **p < 0.01, ***p < 0.001; n.s., not significant.

### NLRP1B and NLRP3 are both required for a full Th1 and Th17 response after *T. mu* colonization

Previous findings demonstrated that host resistance against the pathogenic protist *Toxoplasma gondii* requires the activation of both the NLRP1B and NLRP3 inflammasomes and the innate production of IFN-γ (28-30). Both NLRs get activated through changes in either intra- or extracellular ATP levels and induce the proteolytic processing of pro-IL-1β and pro-IL-18 (31, 32). Given the elevated extracellular concentrations of luminal ATP in *T. mu*^*+*^ mice, we investigated whether *Nlrp1b* or *Nlrp3* contributes to the colonic Th1 and Th17 response following *T. mu* colonization. Analysis of IFN-γ and IL-17A production by Th cells in littermate WT, *Nlrp1b*^*-/-*^, and *Nlrp3*^*-/-*^ mice revealed no significant changes in IFN-γ or IL-17A production during steady state (Figure 3A and B). However, in contrast to their WT counterparts, *Nlrp1b*- and *Nlrp3*-deficient mice failed to show a significant induction of Th1 and Th17 cells in the colonic lamina propria following colonization with *T. mu* (Figure 3A and 3B). Noteworthy, *Nlrp3*^*-/-*^ mice showed a blunted, but significant induction of IL-17^+^IFN-γ^+^ Th cells suggesting a partial requirement for NLRP3 in the regulation Th cell responses following colonization by *T*.*mu* (Figure 3B). *Nlrp1b*^*-/-*^ and *Nlrp3*^*-/-*^ mice did not display increased colonic IL-1β nor IL-18 in the presence of *T. mu*, suggesting a lack of an innate immune response (Supplementary Figure 2A). Accordingly, *T. mu*-colonized *Nlrp1b*^*-/-*^ and *Nlrp3*^*-/-*^ mice lacked both the accumulation of immature Ly6C^+^ macrophages and IL-18R expression on Th1 and Th17 cells (Figure 3C and 3D). These results suggest that the underlying immune adaptation in the colon of *T. mu*-colonized mice requires ATP sensing through NLRP1B and NLRP3.

**Figure 3.**
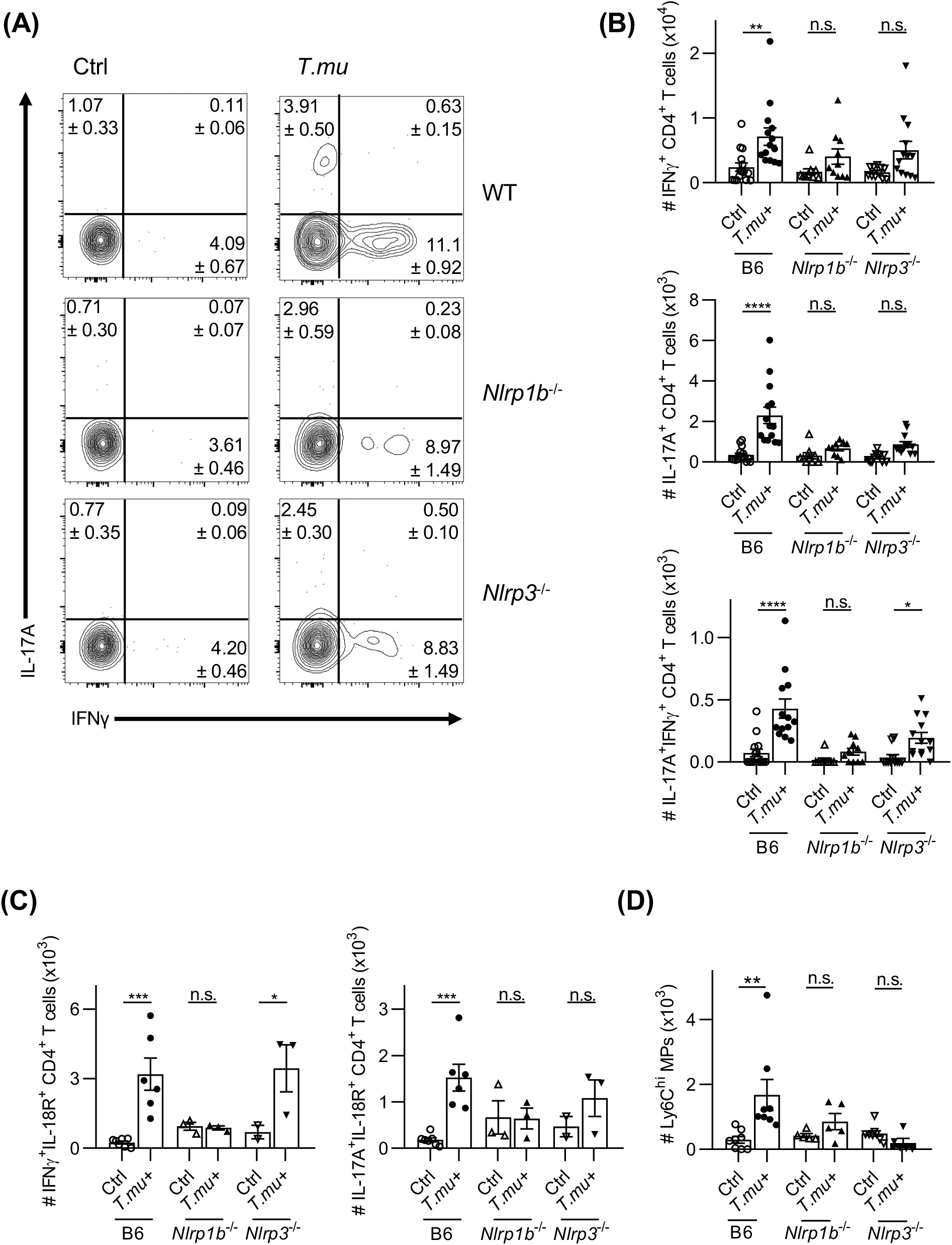
NLRP1B and NLRP3 are required for a full Th1 and Th17 host response after *T. mu* colonization. C57BL/6, *Nlrp1b*^-/-^, or *Nlrp3*^-/-^ mice were either left untreated or orally gavaged with 2 × 10^6^ purified *T. mu*. Colons were harvested 3 weeks later. (A-C) Colonic lamina propria leukocytes were isolated, stimulated for 4 h with PMA and ionomycin in the presence of protein transport inhibitor cocktail, and then stained and analyzed via flow cytometry for (A and B) IFNγ and IL-17A production by CD4^+^ T cells, and (C) IL-18R expression in IFNγ-producing or IL-17A-producing CD4^+^ T cells. (B) Bar graphs show absolute numbers of cell populations represented in respective flow cytometry plots from (A). (D) Colonic lamina propria leukocytes were isolated and analyzed via flow cytometry for Ly6C^hi^ CD64^+^ CD11b^+^ immature macrophages. Data shown is representative of at least three independent experiments with at least three mice per group per experiment. Data show mean ± SEM. Two-way ANOVA with post-hoc Sidak correction was performed; *p < 0.05, **p < 0.01, ***p < 0.001, ****p < 0.0001; n.s., not significant.

### *T. mu* mediates host protection against *Salmonella typhimurium* in an NLRP1B- and NLRP3-dependent fashion

IFN-γ, IL-18, and IL-1β have previously been demonstrated to play a critical role in the host defense against intestinal infection by *Salmonella typhimurium* (*S. typhimurium)* (33, 34). Colonization of WT mice with the commensal protist *T. mu* increased colonic IL-1β levels and led to an accumulation of IL-18R^+^ IFN-γ-producing Th1 cells in an NLRP1B and NLRP3-dependent fashion, implying that the two NLR inflammasomes may be required for the previously shown *T. mu*-induced protection against enteric Salmonella infection (Figure 3) (17). To test this hypothesis, groups of littermate WT, *Nlrp1b*^*-/*^,^*-*^ and *Nlrp3*^*-/-*^ mice were either orally infected with *S. typhimurium* alone, or colonized with *T. mu* for 3 weeks prior to infection. The colonization of mice with *T. mu* did not affect cecal CFUs of *S. typhimurium* across all genotypes, indicating successful infections and the absence of colonization resistance irrespective of NLRP1B, NLRP3, or *T. mu* (Figure 4A). Only littermate control mice carrying *T. mu* failed to show a characteristic loss in cecal weight following infection with *S. typhimurium*, implicating an amelioration of disease in the presence of *T. mu* (Figure 4B). Differences in cecum weights did not significantly change in *Nlrp1b*^*-/-*^ or *Nlrp3*^*-/-*^ mice after *S. typhimurium* infection, even in the presence of *T. mu*, and ranged in between the cecum weights of *T. mu*-free and *T*.*mu*-colonized littermate controls (Figure 4B). Blinded pathology scoring of hematoxylin and eosin-stained Salmonella-infected cecal tissue sections indicated that littermate WT mice colonized with *T. mu* displayed normal histology, with minimal signs of inflammation, in contrast to their *T. mu*-free controls. *Nlrp1b*^*-/-*^ mice presented with comparable levels of tissue damage and inflammatory infiltrates, even in the presence of *T. mu* (Figure 4C and D). While *Nlrp1b*^*-/-*^ mice failed to show protection against Salmonella infection, this effect was less pronounce in *Nlrp3*^*-/-*^ mice. These findings suggest that *T. mu* colonization initiates host resistance to *S. typhimurium* in an NLRP1B-dependent and NLRP3-supported fashion. Damage to the intestinal epithelium following infection facilitates the dissemination of *S. typhimurium* into peripheral organs and results in a breakdown of mucosal barrier function (20). To determine if *Nlrp1b*^*-/-*^ and *Nlrp3*^*-/-*^ mice would display altered dissemination of *S. typhimurium* into peripheral organs in the presence of *T. mu, S. typhimurium* CFUs in the spleen and liver across all experimental groups were quantified. *Nlrp1b*^*-/-*^ and *Nlrp3 /-* mice displayed increased dissemination of *S. typhimurium* into the liver and spleen, even in the presence of *T. mu* (Figure 4E), suggesting that both the NLRP1B and NLRP3 inflammasomes are required for elevated mucosal barrier function driven by the protozoan commensal *T. mu*.

**Figure 4.**
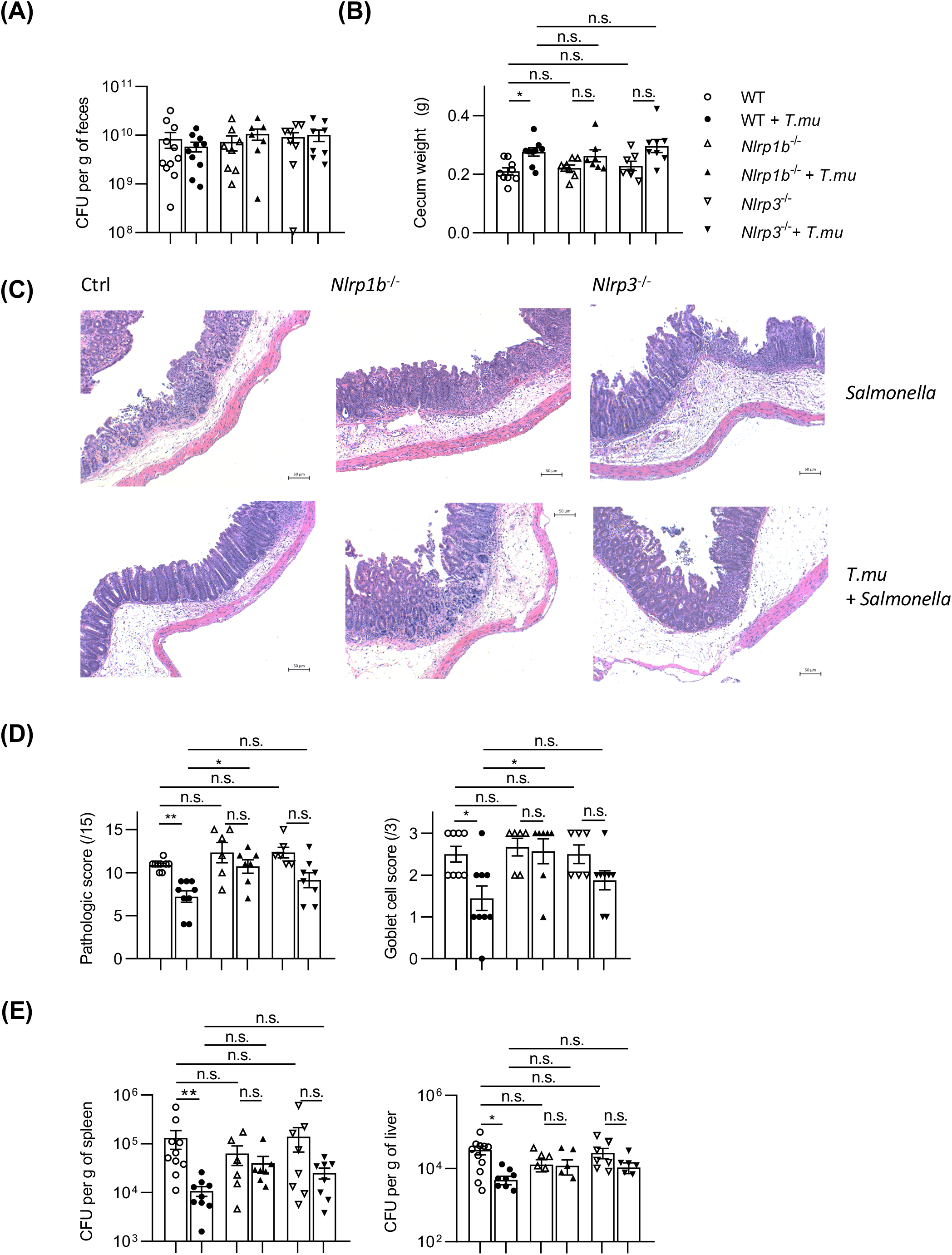
*T. mu* mediates host protection against *Salmonella typhimurium* in an NLRP1B- and NLRP3-dependent fashion. C57BL/6, *Nlrp1b*^-/-^, or *Nlrp3*^-/-^ mice were either left untreated or orally gavaged with 2 × 10^6^ purified *T. mu* 3 weeks prior to *Salmonella typhimurium* infection. (A) Colony-forming units (CFUs) of *S. typhimurium* were measured in fecal pellets 48 h after infection. (B) Cecum weights were measured. (C and D) Blinded pathological scoring was performed on H&E-stained cecal tissue sections. (E) Colony-forming units (CFUs) of *S. typhimurium* were measured in the spleen and liver 48 h after infection. Data shown is representative of at least 3 independent experiments. Data show mean ± SEM. Two-way ANOVA with Sidak correction was performed; *p < 0.05, **p < 0.01, ***p < 0.001, ****p < 0.0001; n.s., not significant.

### Conclusion

In this study, we demonstrate that colonization with the protozoan commensal *T. mu* increases intestinal levels of extracellular ATP. It has previously been demonstrated that some bacteria release ATP into the extracellular space during *in vitro* growth (35). The bacterial microbiome was shown to contribute to the development of Th17 cells and activation of myeloid cells via the release of ATP into the intestinal lumen, suggesting commensal bacteria as a possible source of extracellular ATP (22). Additional sources of extracellular ATP include dying epithelial cells and activated immune cells (21). Colonization with *T. mu* may therefore contribute to the elevated levels of ATP in the extracellular space via direct release of ATP by the protist, induction of epithelial cell death, modulation of the bacterial microbiome, or activation of intestinal immune cells. The possibility of all these sources, individually or in concert, contributing to the elevated levels of extracellular ATP makes this observation a particularly interesting area of future investigation. The resulting alterations in extracellular ATP concentrations following colonization by *T*.*mu* collectively promote the activation of the NLRP1B and NLRP3 inflammasomes.

In addition, we demonstrate that colonization of mice with *T. mu* increases colonic IL-1β levels and induces the accumulation of immature macrophages, IFN-γ-producing Th1 cells, and IL-17A-releasing Th17 cells in the colonic lamina propria. Colonic Th1 and Th17 cells in *T. mu*-colonized mice expressed IL-18R, indicating their ability to respond to IL-18 stimulation. This IL-18R-dependent Th1 and Th17 response required NLRP1B and NLRP3 and may be driven by the inflammasome-dependent accumulation of Ly6C^+^ TNFα-releasing immature macrophages or migratory dendritic cells producing IL-12 (17). In conclusion, the introduction of a protozoan commensal into an established microbiome changes the host immune landscape in the intestine and confers amelioration of Salmonella-induced disease severity. We demonstrate that NLRP1B and in parts NLRP3 are critical in the innate and adaptive immune response following colonization with *T. mu* and are required for the host’s anti-microbial activity against enteric Salmonella infections.

Salmonellosis ranks amongst the world’s top 5 foodborne illness with an annual 1.35M infections, 23.000 hospitalizations and 450 deaths in the United States (ww.cdc.gov). The status of protozoan colonization of those infected is not known. Within the healthy population, ∼10% of individuals are found to carry *T*.*mu*’s closest human relative *Dientamoeba fragilis* (36). Considering observations of Salmonella infections in multiple populations across varying hygiene standards, raises the possibility that underlying commensal protozoan colonization may result in higher asymptomatic Salmonella infections (37). These asymptomatic infections with Salmonella are considered a public health hazard, as spread of infections to non-infected individuals may be promoted more rapidly through these asymptomatic carriers (39). However, the heterogeneity of the infecting Salmonella strains may affect the degree of protection conferred by the protozoan commensal and vice-versa, diversity and heterogeneity of colonizing protozoa may similarly impact protection against enteric infections (38). More research is need to fully grasp the impact of protozoan commensals on host immunity, anti-microbial protection and epidemiology of infectious diseases.

Appreciating the life style of free living wild mice, which often feed on contaminated food, the presence of *Tritrichomonas* spp. as part of their microbiome could present an evolutionary advantage, permitting the consumption of contaminated food that would otherwise result in sickness and disease (Mortha, unpublished observation). Whether protection against other foodborne pathogens may also be regulated by the *T*.*mu-*driven activation of the NLRP1B and NLRP3 inflammasome in mice remains unknown. Our findings warrant considerations for experimental designs in disciplines investigating host-pathogen interactions. Careful surveillance of the intestinal protozoan status in research animals and the implementation of standardized experimental controls are a necessity considering protozoan commensals as member of the healthy mouse microbiome. The observed immune phenotypes in mice carrying *T*.*mu* may result in misinterpretation by investigators if the status of *Tritrichomonas* spp. is not considered, or variable across the studied animal cohort. The horizontal and vertical transmission of *T*.*mu* between mouse lines further calls for the appropriate implementation of littermate controls (40). The methodology used in this study, describes the isolation and colonization of mice with *Tritrichomonas* spp. and provides a guideline for investigators to determine whether their experimental systems may be influenced by a neglected protozoan commensal.

Collectively, we demonstrate that the protozoan commensal *T*.*mu* activates the NLRP1B, and in parts NLRP3 inflammasomes, mediating the remodeling of the host’s intestinal immune landscape to elicit protection against the pathology caused by enteric Salmonella infections. Future studies will investigate the protective role of *T*.*mu* against infections with other foodborne pathogens.

## Disclosures

The authors have no financial conflicts of interest.

## Acknowledgments

We thank members of the #onlylabever for their discussion and experimental assistance. We acknowledge Dr. Nathalie Simard and Janine Charron (Temerty Faculty of Medicine Flow Cytometry Core) for providing cell-sorting support. We thank Dr. Dana Philpott and Dr. Juan-Carlos Zuniga-Plücker for critical reading of the manuscript.

## Footnotes

### Grant support

P.C. is supported by an Ontario Trillium Scholarship and a Vanier/NSERC Canada Graduate Scholarship. K.B. is supported by a Banting Postdoctoral Fellowship. L.N. is supported by an Ontario Graduate Scholarship. A.M. is supported by a CIHR-Project Grant (388337) and a NSERC-Discovery Grant (RGPIN-2019-04521). A.M. is the Tier 2 Canadian Research Chair in Mucosal Immunology and supported by the Tier 2 CRC-CIHR program.

## Abbreviations used in this article

*T. mu*: *Tritrichomonas musculis*
ATP: adenosine 3’-triphosphate
WT: wild-type
TLR: Toll-like receptor
NLR: Nod-like receptor

## Figure legends

**Supplementary Figure 1.**
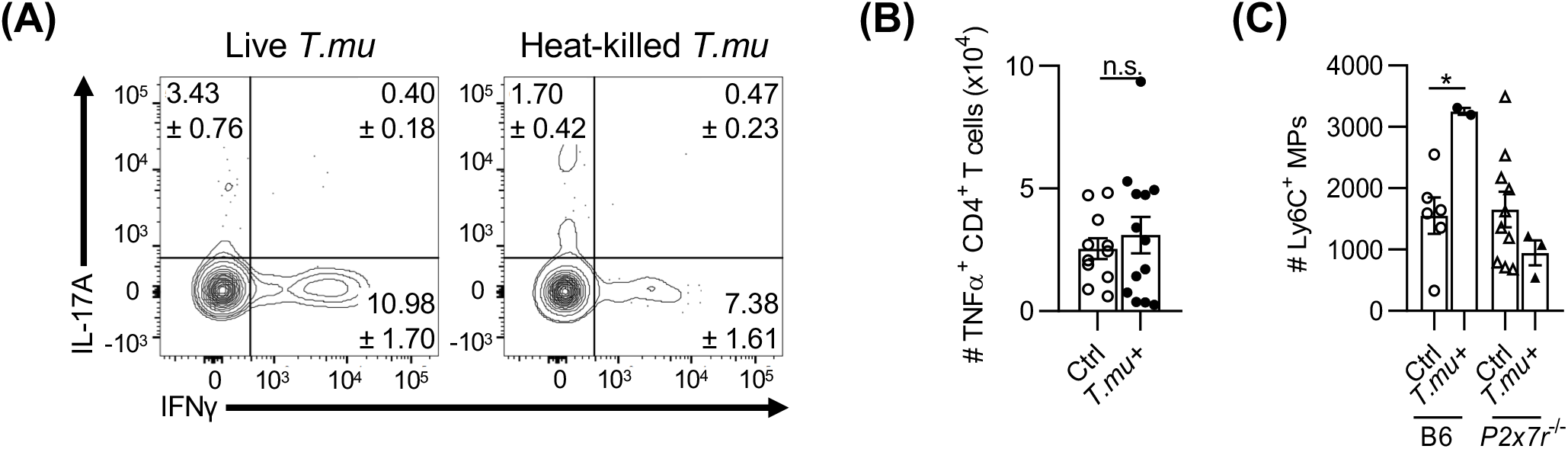
Viable *T. mu* drives Th cell activation but does not alter TNFα production in Th cells. (A) 2 × 10^6^ purified *T. mu* was either immediately orally gavaged or incubated at 95°C for 1 min prior to gavage into C57BL/6 mice. 3 weeks later, colonic lamina propria leukocytes were isolated, stimulated for 4 h with PMA and ionomycin in the presence of protein transport inhibitor cocktail, and then stained and analyzed via flow cytometry for IFN-γ and IL-17A production by CD4^+^ T cells. (B) C57BL/6 mice were either left untreated or orally gavaged with 2 × 10^6^ purified *T. mu*. 3 weeks later, colonic lamina propria leukocytes were isolated, stimulated for 4 h with PMA and ionomycin in the presence of protein transport inhibitor cocktail, and then stained and analyzed via flow cytometry for TNFα production by CD4^+^ T cells. (C) C57BL/6 or *P2x7r*^-/-^ mice were either left untreated or orally gavaged with 2 × 10^6^ purified *T. mu*. 3 weeks later, colonic lamina propria leukocytes were isolated and analyzed via flow cytometry for Ly6C^hi^ CD64^+^ CD11b^+^ immature macrophages. Data shown is representative of at least three independent experiments. Data show mean ± SEM. Student’s t test was performed for (B), two-way ANOVA with post-hoc Sidak correction was performed for (C); *p < 0.05, **p < 0.01, ***p < 0.001; n.s., not significant.

**Supplementary Figure 2.**
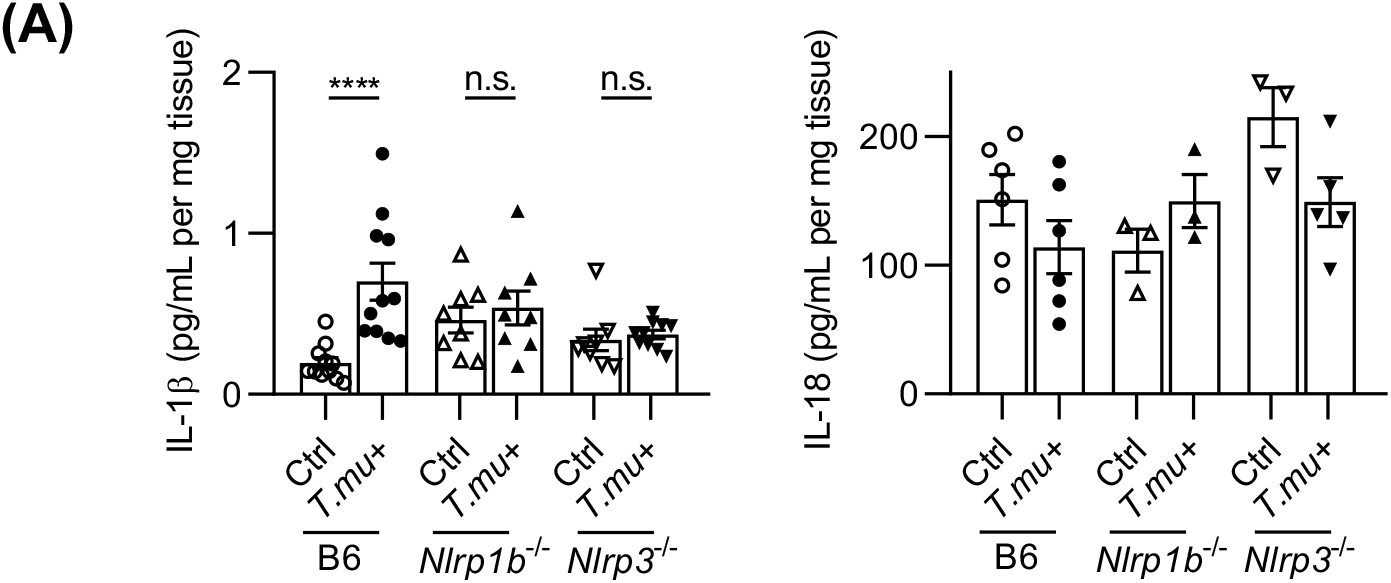
NLRP1B- and NLRP3-dependent release of IL-1β following colonization with *T. mu*. C57BL/6, *Nlrp1b*^-/-^, and *Nlrp3*^-/-^ mice were either left untreated or orally gavaged with 2 × 10^6^ purified *T. mu*. 3 weeks later, colonic explants were placed in RPMI + gentamicin for 30 min, then incubated overnight at 37C in 500 uL RPMI supplemented with gentamicin, penicillin/streptomycin, and 5% FBS. Supernatants were collected for IL-1β and IL-18 ELISA analysis. Data show mean ± SEM. Two-way ANOVA with post-hoc Sidak correction was performed; *p < 0.05, **p < 0.01, ***p < 0.001; n.s., not significant.

## Notes

### Competing Interest Statement

The authors have declared no competing interest.

